# Human Herpes Virus 6 (HHV-6) - Pathogen or Passenger? A pilot study of clinical laboratory data and next generation sequencing

**DOI:** 10.1101/236083

**Authors:** Colin Sharp, Tanya Golubchik, William F. Gregory, Anna L. McNaughton, Nicholas Gow, Mathyruban Selvaratnam, Alina Mirea, Dona Foster, Monique Andersson, Paul Klenerman, Katie Jeffery, Philippa C. Matthews

**Affiliations:** The Roslin Institute, University of Edinburgh, Easter Bush, Midlothian, Edinburgh, Scotland, UK; Edinburgh Genomics, Ashworth Laboratories, University of Edinburgh, Edinburgh, EH9 3FL, Scotland, UK; The Wellcome Trust Centre for Human Genetics, Roosevelt Drive, Oxford, OX3 7BN, UK; Big Data Institute, University of Oxford, Old Road, Oxford, OX3 7FZ UK; Nuffield Department of Medicine, Peter Medawar Building for Pathogen Research, University of Oxford, South Parks Road, Oxford, OX1 3SY, UK; Department of Infectious Diseases and Microbiology, Oxford University Hospitals NHS Foundation Trust, John Radcliffe Hospital, Headley Way, Headington, Oxford, OX3 9DU, UK; NIHR Biomedical Research Centre, Nuffield Department of Medicine, University of Oxford, John Radcliffe Hospital, Headley Way, Headington, Oxford, OX3 9DU, UK

**Keywords:** Human herpesvirus 6, diagnosis, encephalitis, haematology, metagenomics, next generation sequencing

## Abstract

**Background:** Human herpes virus 6 (HHV-6) is a ubiquitous organism that can cause a variety of clinical syndromes ranging from short-lived rash and fever through to life-threatening encephalitis.

**Objectives:** We set out to generate observational data regarding the epidemiology of HHV-6 infection in clinical samples from a UK teaching hospital and to compare different diagnostic approaches.

**Study design:** First, we scrutinized HHV-6 detection in samples submitted to our hospital laboratory through routine diagnostic pathways. Second, we undertook a pilot study using Illumina next generation sequencing (NGS) to determine the frequency of HHV-6 in CSF and respiratory samples that were initially submitted to the laboratory for other diagnostic tests.

**Results:** Of 72 samples tested for HHV-6 by PCR at the request of a clinician, 24 (33%) were positive for HHV-6. The majority of these patients were under the care of the haematology team (30/41, 73%), and there was a borderline association between HHV-6 detection and both Graft versus Host Disease (GvHD) and Central nervous system (CNS) disease (p=0.05 in each case). We confirmed detection of HHV-6 DNA using NGS in 4/20 (20%) CSF and respiratory samples.

**Conclusions:** HHV-6 is common in clinical samples submitted from a high-risk haematology population, and enhanced screening of this group should be considered. NGS can be used to identify HHV-6 from a complex microbiomee, but further controls are required to define the sensitivity and specificity, and to correlate these results with clinical disease. Our results underpin ongoing efforts to develop NGS technology for viral diagnostics.

## ABBREVIATIONS

cDNA: complementary DNA
CiHHV-6: Chromosomally integrated HHV-6
CMV: Cytomegalovirus
CNS: central nervous system
CSF: cerebrospinal fluid
EPR: Electronic patient record
GvHD: graft versus host disease
HHV-6: Human herpes-virus-6
HLA: human leucocyte antigen
HSCT: Haematopoietic stem cell transplant
HSV: Herpes Simplex Virus
HTLV: Human T cell lymphotrophic virus
NGS: Next Generation Sequencing
OUH: Oxford University Hospitals NHS Foundation Trust
PCR: Polymerase Chain Reaction
RSV: Respiratory Syncytial Virus
VZV: Varicella Zoster Virus

## BACKGROUND

Human herpes-virus-6 (HHV-6) is a human beta-herpesvirus (1). Like its close relative human cytomegalovirus (CMV), it is ubiquitous, has the potential for latency followed by chronic low level replication or reactivation, and may modulate immune responses to other pathogens (2-5). In children, is usually asymptomatic or associated with self-limiting fever and rash. It is also associated with encephalitis either as a primary agent (6) or as a result of reactivation in the setting of encephalitis / meningitis caused by other pathogens, in which context it appears generally benign (7). At the other end of the spectrum, HHV-6 can reactivate in the context of severe sepsis (8) and is a cause of potentially life-threatening pathology in patients with haematological malignancy, usually following HSCT (haematopoietic stem cell transplant) (9-15).

HHV-6 variants A and B share approximately 90% homology (16). HHV-6A accounts for more CiHHV-6 (17), while HHV-6B is associated with acute infection, childhood rash/fever (4), and reactivation following HSCT (16). Although HHV-6A is less common in CNS disease (18), it may be more aggressive when present (16). HHV-6 transmission is predominantly via respiratory secretions or saliva, but can also be vertical as a result of chromosomally-integrated HHV-6, ‘CiHHV-6’ (17, 19), leading either to episodic reactivation (17), or to persistent viraemia (typically ≥5.5 log10 copies/ml (˜300,000 copies/ml) in blood (20)).

HHV-6 laboratory diagnostics raise a number of challenges: what sample type to test, in which patient groups to focus, and how to interpret a positive test result. There is increasing interest in ‘next generation sequencing’ (NGS) approaches to diagnosis of many pathogens (21-25) but optimization is required for these pipelines, including tackling high proportions of human reads, differentiating between pathogenic organisms and commensal / environmental flora, and determining thresholds at which the identification of an organism is likely to be clinically significant (26).

## OBJECTIVES

We set out to review HHV-6 data from the local diagnostic microbiology laboratory, to determine the patterns of clinical testing for HHV-6. Second, we screened randomly selected CSF and respiratory samples using PCR and NGS in a small pilot study to ascertain the extent to which this virus can be detected in routine samples. Together, these aim to describe the distribution of HHV-6 in local clinical samples, and to evaluate the contribution made by different diagnostic tools, thereby informing ongoing development of laboratory protocols for diagnosis.

## STUDY DESIGN

### Study site, cohorts and ethics

Clinical data and samples from between 2013-2016 were collected from the microbiology department at Oxford University Hospitals (OUH) NHS Foundation Trust, a large tertiary referral teaching centre in the UK (http://www.ouh.nhs.uk/). This study pertains to the analysis of two separate sample cohorts (Suppl data set 1):

i. Samples submitted to the laboratory by clinicians with a request for **HHV-6** screening (ID numbers prefixed **HHV**). This is undertaken at the request of the clinical team when deemed clinically relevant;
ii. Samples which had no request for HHV-6 testing, but had completed routine diagnostic laboratory testing for other indications and were used for **v**iral **s**equencing studies (ID numbers prefixed **VS**).

Approval for retrospective collection of clinical and laboratory data was granted by the OUH Clinical Audit Committee (HHV cohort). Testing of consecutive anonymised laboratory samples (VS cohort) was approved by local Research Services and through review via the UK Integrated Research Application System (REC Reference 14/LO/1077).

### Collection of local laboratory data (HHV cohort)

We undertook an electronic search of the OUH Microbiology laboratory system to identify all instances of an HHV-6 test (antibody or viral load) being requested over three years commencing 1-Jan-2013, and recorded age, sex, sample type and patient location. We used the Electronic Patient Record (EPR) to determine underlying diagnosis. Follow-up data for survivors were available for a median of 25 months (range 316 – 1374 days). Diagnostic tests were undertaken by in-house real time PCR assays (27) at two different National Reference Laboratories (Colindale up to 1-Oct-2014, and subsequently Bristol).

### Data collection and statistical analysis

We used GraphPad Prism v.6.0f for statistical analysis with Fisher’s Exact Test to identify differences between binary groups, and the Mann Whitney U test for continuous variables. Multivariate regression analysis was undertaken using open access on-line software (https://docs.google.com/spreadsheets/u/0/).

### Testing CSF and respiratory samples (VS cohort)

We identified 100 CSF samples and 100 respiratory samples (throat swabs (n=22), nasopharyngeal aspirates (n=42), endotracheal aspirates (n=4), and bronchoalveolar lavage samples (n=32)) representing a ‘high risk’ subgroup based on the following criteria:

i. For CSF, the clinical request for testing included viral causes of meningitis and encephalitis;
ii. For respiratory samples, the patient had a clinical history of immunocompromise and/or was on the intensive care unit.

### PCR and NGS (VS cohort)

Samples had all undergone clinical laboratory testing and were stored at −80°C prior to further processing. We selected consecutive samples; the only exclusion criterion was inadequate sample volume (<200 ul). For each sample, we documented patient age group and clinical location, and recorded the clinical information supplied with the sample and routine microbiology laboratory data.

Nucleic acids were extracted from 200μl of each sample using the AllPrep DNA/RNA Mini Kit (Qiagen) and recovered in 30μl of nuclease-free water. To allow for broad detection of HHV-6 and other known herpesviruses, 4ul of DNA was used as template for a consensus PCR primer as previously published (28). The results were visualised on 2% agarose gels and amplicons from positive reactions were cut from the gels for sequencing. Direct amplicon sequencing was performed using BigDye Terminator v3.1 (Applied Biosystems) according to the manufacturer’s instructions with both second round primers. Sequencing reactions were read by Edinburgh Genomics and assembled using SSE v1.2 (29).

We selected a random subset of 20 samples for Illumina sequencing. Methods are described in detail in another manuscript (30).

## RESULTS

### Routine clinical samples received for HHV-6 testing (HHV cohort)

During the three-year study period, our clinical laboratory received 85 samples for HHV-6 testing (Fig 1A; suppl data set 1). In total, 41 patients were tested; 22 M:19 F; median age 52 years (range 2-71; IQR 39-61). Central nervous system (CNS) disease (encephalitis, encephalopathy, meningism, seizures) was present in 9/41 (22%). Most patients were under haematology care (30/41; 73%), of these, 5/30 (17%) had Graft versus Host Disease (GvHD). 72/85 samples were tested for HHV-6 DNA by PCR at the reference laboratory; 24/72 (33%) were positive. Among individual patients tested, 15/41 (37%) were PCR positive at ≥1 timepoint (Fig 1A). Of seven samples that were subtyped, all were HHV-6B.

**Figure 1:**
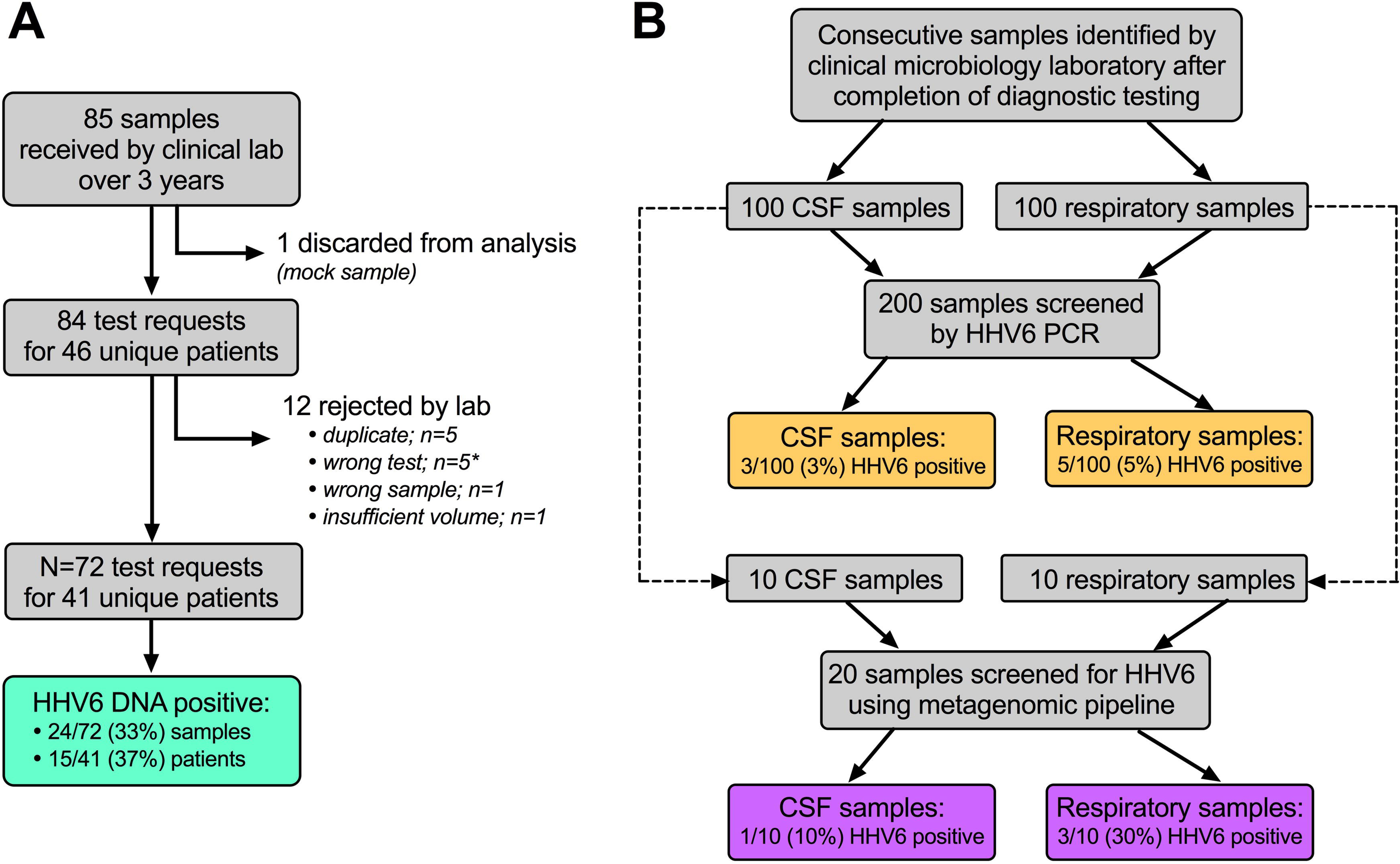
Schematic summary of work flow and output of HHV-6 testing among clinical samples from a UK teaching hospital. A: Flow diagram showing number of samples submitted to a hospital diagnostic microbiology laboratory with a clinical request for HHV-6 testing (sample ID’s prefixed with HHV). *’Wrong test’ indicates request for HHV-6 antibody (rather than PCR). B: Flow diagram showing consecutive random samples (sample ID’s prefixed with VS) screened for herpesviruses (i) by PCR with HHV-6 confirmed by sequencing, and (ii) by a metagenomic approach. These samples had been submitted to the clinical laboratory for other reasons, and had reached the end of their diagnostic testing pathway.

Positive HHV-6 status was not statistically associated with age, sex, or haematological malignancy (Table 1). There was a borderline association with both GvHD and CNS disease (both p=0.05; Table 1; Fig. 2A and B), but this is difficult to interpret as patients in these groups are more likely to be selected for HHV6 testing. On multiple logistic regression analysis, there was no relationship between HHV-6-positivity and any other characteristic (Table 1).

**Figure 2:**
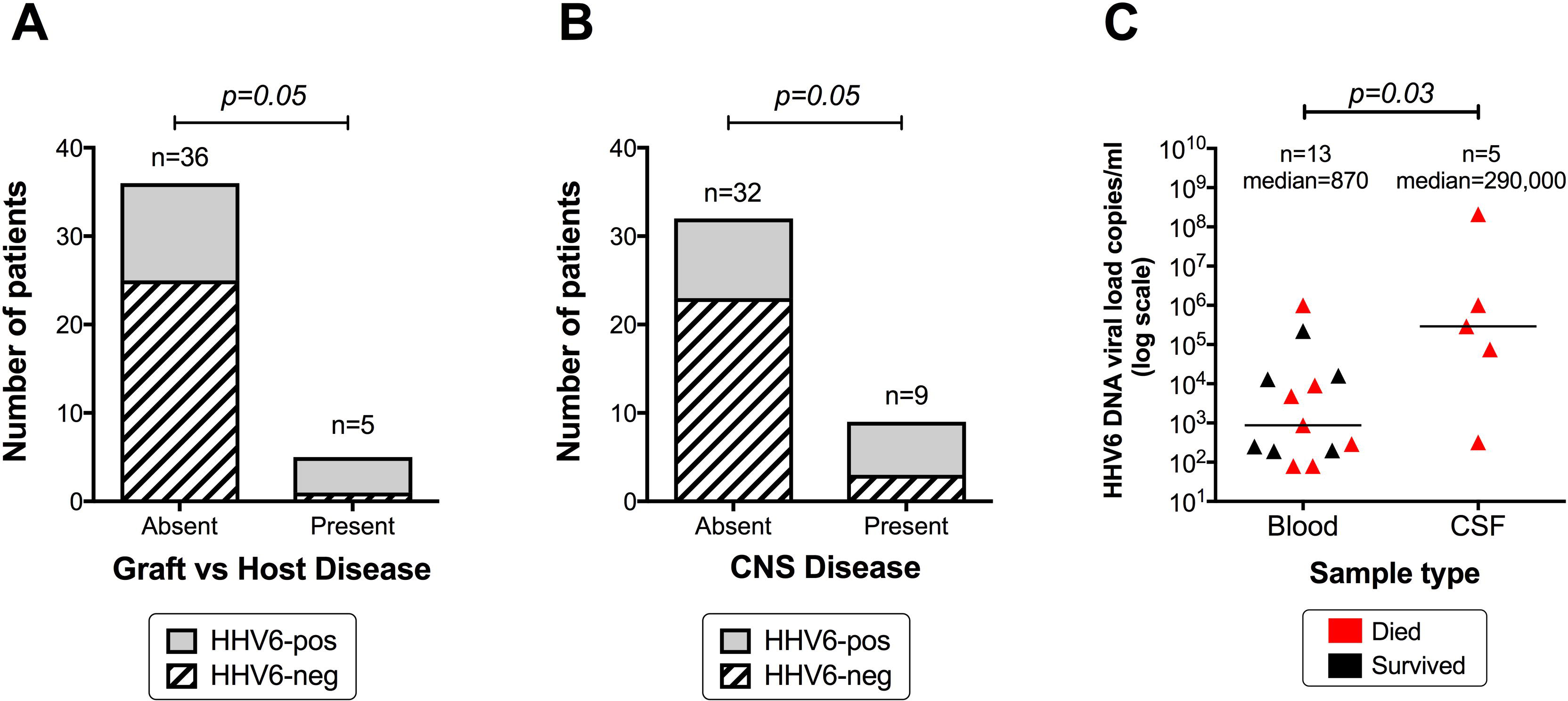
Results of clinical laboratory testing for HHV-6 in blood and CSF samples at the request of the clinical team. A: Relationship between presence or absence of Graft vs Host Disease (GvHD) and HHV-6 status; P value by Fisher’s exact test. B: Relationship between presence or absence of a clinical Central Nervous System (CNS) syndrome and HHV-6 status; P value by Fisher’s exact test. C: HHV-6 viral loads in blood and CSF; P value by Mann Whitney test; individuals who died are shown in red. In all three panels, the numbers at the top of each column show the total number of patients represented.

**Table 1:** Relationship between HHV-6 status and clinical/demographic features in 41 patients with samples submitted for HHV-6 testing to a clinical UK microbiology lab.

|  | HHV-6 PCR<br>positive<sup>a</sup><br>(n=15<br>patients) | HHV-6 PCR<br>negative<br>(n=26<br>patients) | p-value<sup>b</sup><br>(univariate<br>analysis by<br>Fisher's<br>Exact test) | p-value<sup>b</sup><br>(multivariate<br>analysis by<br>logistic<br>regression) |
| --- | --- | --- | --- | --- |
| **Sample type<sup>c</sup>:**<br>**blood** | 9/15<br>(60%) | 15/26<br>(58%) | 1.0 | 0.7 |
| **Sample type<sup>c</sup>:**<br>**CSF** | 3/15<br>(20%) | 4/26<br>(15%) | 0.7 | 0.8 |
| **Age ≥50 years** | 10/15<br>(67%) | 13/26<br>(50%) | 0.3 | 0.3 |
| **Male sex** | 8/15<br>(53%) | 14/26<br>(54%) | 1.0 | 0.2 |
| **Haematological**<br>**malignancy**<br>**present** | 13/15<br>(87%) | 17/26<br>(65%) | 0.2 | 0.9 |
| **Graft versus**<br>**host disease**<br>**present** | 4/15<br>(27%) | 1/26<br>(4%) | **0.05** | 0.1 |
| **CNS<sup>d</sup> disease**<br>**present** | 6/15<br>(40%) | 3/26<br>(12%) | **0.05** | 0.1 |
| **Solid organ**<br>**transplant**<br>**present** | 0/15<br>(0%) | 2/26<br>(8%) | 0.5 | 1.0 |
| **Death<sup>e</sup>** | 11/15<br>(73%) | 11/25<br>(44%) | 0.1 | 0.2 |

<sup>a</sup> In the case of patients tested more than once, any instance of PCR positive is recorded as positive; <sup>b</sup> Sample types were not documented for 3 patients testing PCR-positive, and for 7 patients testing PCR-negative; <sup>c</sup> CNS=central nervous system; <sup>d</sup> Prospective survival data were available for 40/41 patients.

In this cohort, 23/41 (56%) of patients died, at a median age of 56 years. Among these, 11/23 (48%) had tested HHV-6-PCR positive, compared to 4/18 (22%) of surviving patients (p=0.1). All those who died did so within 19 months of the HHV-6 test (range 1-552 days, median 163 days), and all patients with HHV6 detected in the CSF died (Fig 2C). We did not have sufficient clinical data to determine cause of death, and HHV6 may be a bystander in this complex cohort.

### Quantification of HHV-6 viral load in blood and CSF (HHV cohort)

HHV-6 DNA was quantified in 23 samples from ten patients (Fig 2C). The levels varied from below the threshold for accurate quantification, to patient HHV-012 with >1.0×10^6^ copies/ml in blood and 2.0×10^8^ DNA copies/ml in CSF, suggesting CiHHV-6. Patient HHV-007, an adult with haematological malignancy, had HHV-6 DNA detected in both blood and CSF (9×10^4^ DNA copies/ml vs. 3×10^5^ copies/ml, respectively). This patient had limbic encephalitis, and the raised HHV-6 titre in CSF compared to blood is in keeping with a localized CNS pathology caused by the virus.

Among a total of 72 tests, 46 were longitudinal samples from thirteen individual patients. There was no statistical association between multiple HHV-6 tests and other factors that might predict disease severity (p>0.1 for age, sex, mortality, haematological malignancy, transplant, intensive care location, documented co-infection in multiple regression analysis).

### Screening laboratory samples for HHV-6 by PCR (VS cohort)

We screened CSF (n=100) and respiratory (n=100) samples for herpesvirus DNA by PCR and sequencing (Fig 1B; Suppl data set 1), identifying HHV-6 in 3/100 CSF samples and 5/100 respiratory samples (Table 2). Four of the eight positive cases were age <5 years, representing the age group in whom primary HHV-6 infection is most likely. However, in four samples (two adults and two children) an alternative pathogen was identified by the clinical lab (Table 2), illustrating the difficulty in distinguishing between HHV-6 as a primary pathogen, a co-infecting agent contributing to pathology, or an innocent bystander.

**Table 2:** Summary of eight clinical samples (from a total of 200 consecutive samples selected for screening) that tested positive for HHV-6 DNA by PCR.

| Study ID number | Sample type | Age group (years) | Location | Clinical details | CSF cell count: total WCC (cells/μl) | CSF cell count: lymphs (cells/μl) | CSF cell count: RBC (cells/μl) | Bacterial culture results from routine clinical lab | Molecular diagnostics (PCR) from routine clinical lab |
| --- | --- | --- | --- | --- | --- | --- | --- | --- | --- |
| VS013 | CSF | 20-24 | Emergency Department | Headache, ? SAH | 346 | 342 | 70 | No growth | VZV detected |
| VS014 | CSF | <5 | Paediatric ICU | ? Meningitis | 8 | 6 | 8 | No growth | No virus detected |
| VS022 | CSF | 60-64 | Medicine | Confusion, temporal attenuation on CT brain | 28 | 26 | 28 | No growth | HSV type 1 detected |
| VS114 | NPA | <5 | Paediatric ICU | ? Bronchiolitis | n/a | n/a | n/a | Not requested | No organism detected |
| VS121 | NPA | <5 | Paediatric ICU | LRTI | n/a | n/a | n/a | Not requested | Rhinovirus detected |
| VS127 | BAL | 65-69 | Chest Unit | Pneumonia | n/a | n/a | n/a | No growth | No organism detected |
| VS152 | BAL | 55-59 | Chest Unit | ? TB | n/a | n/a | n/a | *Streptococcus viridans* | No organism detected |
| VS183 | NPA | <5 | Paediatrics | Viral infection | n/a | n/a | n/a | Not requested | Meta-pneumovirus detected |

CSF=cerebrospinal fluid, NPA=nasopharyngeal aspirate, BAL=broncho-alveolar lavage, HSV = Herpes Simplex Virus, ICU=intensive care unit, SAH=subarachnoid haemorrhage, CT=computer tomography, LRTI=lower respiratory tract infection, TB=tuberculosis, PCR=polymerase chain reaction, VZV=varicella zoster virus. '?' in clinical details column reflects potential but unconfirmed diagnosis being raised as a query by the requesting clinician.

The overall HHV-6 prevalence of 4% in this random group of samples is significantly lower than the 33% rate obtained from the samples in which the clinician had requested HHV-6 testing (p<0.0001; Fisher’s Exact Test). This suggests that the targeting of the highest risk groups (primarily patients with haematological malignancy) for HHV-6 testing is appropriate, and that the high prevalence of detectable HHV-6 DNA in this group is not merely reflective of universal reactivation of herpesviruses in a hospital cohort.

### Screening laboratory samples for HHV-6 DNA by NGS (VS cohort)

We screened a subset of 20 samples (10 CSF and 10 respiratory; Fig 1B) using a metagenomic (NGS) approach (30), identifying HHV-6 in four samples that we deemed ‘positive’; one CSF and three respiratory samples (Table 3; Fig 3). The number and distribution of reads in each sample is shown in Fig 4A. Sample VS183 was designated HHV-6B by Kraken, while the other three positive samples were identified as HHV-6A (Fig 4B). The coverage of the HHV-6 genome was incomplete, but multiple reads distributed across the genome (Fig 4B), add confidence to the conclusion that HHV-6 DNA is genuinely present in these samples. The sequence data have been uploaded to European Nucleotide Archive (ENA) https://www.ebi.ac.uk/ena; HHV6 sequence accession numbers ERS1980462 (sample ID VS067), ERS1980463 (VS183), ERS1980464 (VS200) and ERS1980465 (VS207). Links to the full metagenomic sequence set can be found in our supporting Data Note (30).

**Figure 3:**
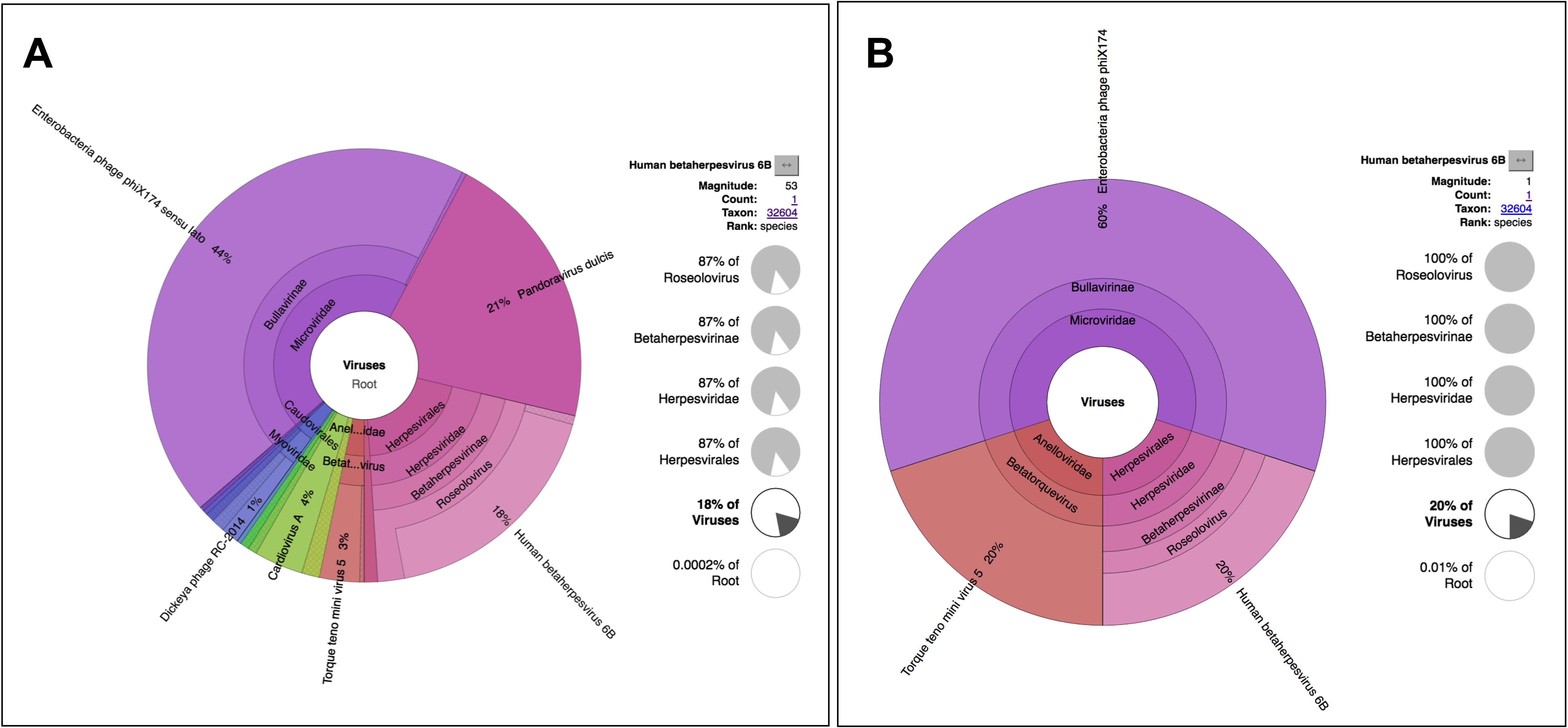
Multi-layered pie charts generated to visualize the metavirome from a respiratory sample taken from a child. Krona was used to generate the metagenomic visualization of these data (35). The sample was a nasopharyngeal aspirate taken from patient ID VS183 (a child age <5 with a clinical syndrome described by the requesting clinician as ‘viral infection’). (A) HHV-6B (in pink) shown as a proportion of all viral reads; (B) HHV-6B (in pink) shown as a proportion of all virus contigs. The other two predominant viruses represented in both panels are Torque Teno Mini Virus (in red), a ubiquitous and non-pathogenic virus, and Enterobacteria phage Phi X (in purple), which is an artefact of the sequencing method (spike used for positive control in Illumina sequencing run); these illustrate a high proportion of sequence reads generated from organisms that are not clinically significant.

**Figure 4:**
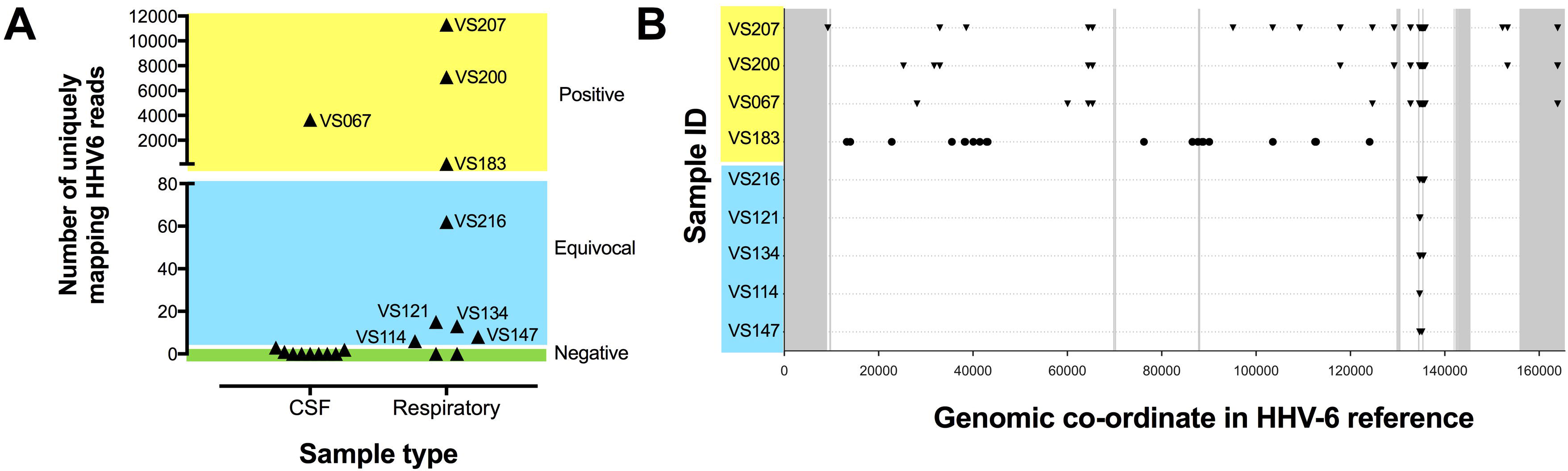
Detection of HHV-6 DNA in unselected CSF and respiratory samples. A: Read numbers determined by next generation sequencing (NGS) according to sample type. The y-axis refers to the number of uniquely mapping HHV6 reads, representing deduplicated read numbers (Q>30). The areas shaded green, blue and yellow represent suggested thresholds for samples to be classified as negative, equivocal or positive, respectively. White gaps between the colours illustrate that the exact position of the boundaries between these areas are uncertain. Those classified as equivocal were all respiratory samples, and the sequences clustered in the repeat region of the genome, suggesting lower specificity for HHV-6. B: Plots generated with Burrows-Wheeler Aligner (BWA) to illustrate coverage of HHV-6 genomes against consensus. The x-axis of the plot represents the full length genome for HHV-6, with HHV-6A shown in triangles, and HHV-6B shown in circles. From a total of 20 samples tested, we show data for samples from which we generated any HHV-6 reads. The beginning position of each read is indicated. The grey bars indicate repeat regions (low variability) as defined in the methods; for a sample to be considered positive, we stipulated that the reads should fall outside these regions of the genome.

**Table 3:** Summary of HHV-6 results from twenty clinical samples screened for pathogens by a metagenomic (NGS) approach.

| Study ID number | Sample type | Age (years) | Clinical details on request form | CSF Lymphocyte count (cells/μl) | Clinical laboratory virology result | HHV-6 PCR result | HHV-6 detection by NGS approach |
| --- | --- | --- | --- | --- | --- | --- | --- |
| VS011 | CSF | <5 | Fever | 2 | No virus detected | Negative | Negative |
| VS012 | CSF | <5 | VP shunt | 4 | No virus detected | Negative | Negative |
| VS033 | CSF | 60-64 | No data | 0 | No virus detected | Negative | Negative |
| VS036 | CSF | <5 | No data | 0 | No virus detected | Negative | Negative |
| VS047 | CSF | 5-9 | No data | 0 | No virus detected | Negative | Negative |
| VS060 | CSF | 40-44 | Previous high CSF white cell count (50). | 32 | No virus detected | Negative | Negative |
| VS067 | CSF | 50-54 | Confusion, headache, malaise, fever | 438 | No virus detected | Negative | Positive (BWA HHV-6A, 3653 reads, 1% coverage) |
| VS071 | CSF | 70-74 | Headache, fever, cognitive impairment | 0 | No virus detected | Negative | Negative |
| VS079 | CSF | 40-44 | Sepsis, Confusion, Encephalopathy | 2 | No virus detected | Negative | Negative |
| VS088 | CSF | 20-24 | Mixed sensory and motor symptoms | 0 | No virus detected | Negative | Negative |
| VS114 | Resp (NPA) | 75-79 | Possible bronchiolitis | N/A | No organism detected | Positive | Equivocal (HHV-6A; BWA mapping within repeat region only) |
| VS121 | Resp (NPA) | <5 | Lower respiratory tract infection | N/A | Rhinovirus detected | Positive | Equivocal (HHV-6A; BWA mapping within repeat region only) |
| VS134 | Resp (BAL) | 55-59 | Neutropenic; sudden respiratory deterioration | N/A | No organism detected | Negative | Equivocal (HHV-6A; BWA mapping within repeat region only) |
| VS147 | Resp (BAL) | 50-54 | ?TB | N/A | No organism detected | Negative | Equivocal (HHV-6A; BWA mapping within repeat region only) |
| VS167 | Resp (NPA) | <5 | Tracheostomy, NIV, increasing secretions | N/A | Rhinovirus detected | Negative | Negative |
| VS177 | Resp (NPA) | <5 | LRTI, CXR changes | N/A | Meta-pneumo-virus detected | Negative | Negative |
| VS183 | Resp (NPA) | <5 | Viral infection | N/A | Meta-pneumo- | Positive | Positive (HHV-6B; 93 reads; |

|  |  |  |  |  | virus detected |  | BWA 1% coverage) |
| --- | --- | --- | --- | --- | --- | --- | --- |
| VS200 | Resp (NPA) | <5 | DKA, increased respiratory secretions, | N/A | RSV detected | Negative | Positive (HHV-6A; 7077 reads; BWA 1% coverage) |
| VS207 | Resp (BAL) | <5 | Respiratory arrest | N/A | Rhinovirus detected | Negative | Positive (HHV-6A; 11305 reads; BWA 1% coverage) |
| VS216 | Resp (NPA) | <5 | LRTI,? TB | N/A | No organism detected | Negative | Equivocal (HHV-6A; BWA mapping within repeat region only) |

BAL = broncho-alveolar lavage; BWA = Burrows-Wheeler Aligner (33); DKA = diabetic ketoacidosis; LRTI = lower respiratory tract infection; NIV = non invasive ventilation; NGS = next generation sequencing; NPA = nasopharyngeal aspirate; N/A = not applicable; TB = tuberculosis; VP = ventriculo-peritoneal. Read counts and thresholds for positivity are shown in Fig 4.

In VS067, the clinical syndrome was not explained by other diagnostic results, and HHV-6 infection was a plausible agent of the clinical syndrome (meningoencephalitis). In the three other cases, other primary pathogens had been identified (Table 3), although it is plausible that HHV-6 could have been a contributory agent.

Among the total of eight samples that were HHV-6 positive by conventional PCR, three were tested on the NGS pipeline (samples VS114, VS121 and VS183). HHV-6 reads were detected by NGS in all three, although two had low numbers of reads leading us to classify them as equivocal (Fig 4A,B). Conversely, in four samples where HHV-6 was detected by NGS, only one of these was positive by PCR.

## DISCUSSION

This preliminary study provides insight into the distribution of HHV-6 in a range of samples in a UK teaching hospital, with the long-term aim of informing improvements in diagnostic testing. Although our NGS data only represent a small pilot study, we have identified only a single other reference to date that describes a metagenomic approach to the diagnosis of HHV-6 infection (6). Although HHV-6 can reactivate in the context of any critical illness (8), we here found a higher prevalence in clinical samples taken from a high-risk group (HHV cohort) than in laboratory samples representing an unwell hospital population (VS cohort), suggesting that HHV-6 is not simply reactivating across the board in hospitalized patients. The combination of a high rate of HHV-6 DNA detection and the high mortality in the clinical (HHV) cohort, suggest that we should consider lowering our threshold for testing in this context. Longitudinal HHV-6 PCR testing should generally be reserved for monitoring response to therapy, but if a high index of suspicion exists (e.g. a profoundly immunosuppressed patient who becomes encephalopathic in the absence of an alternative explanation), then serial testing may be helpful.

This study was not designed to evaluate the use or outcome of antiviral therapy as this information could not robustly be captured retrospectively. There are some data to suggest good outcomes from treatment of symptomatic viraemia (31), with ganciclovir and/or foscarnet. However, there is no universally agreed definition of clinical disease or threshold for therapy and this area is not well informed by clinical trials (32); the gains made by testing early have to be carefully balanced against the possibility of identifying patients with bystander viral reactivation in whom the toxicity and side-effects of treatment would not be justified. Further studies, including NGS, may determine whether different clinical syndromes are consistently associated with the two HHV-6 subtypes.

For samples with low HHV-6 copy numbers, PCR is anticipated to be more sensitive than NGS; this is illustrated by two samples that tested positive by PCR but were equivocal by NGS. However, the overall proportion of samples testing positive for HHV-6 DNA was higher by NGS than by PCR. This may at least in part be accounted for by PCR using highly degenerate herpesvirus primers, followed by sequencing primers that have a higher degree of sequence homology for HHV-6B than HHV-6A (28).

In screening by NGS, we identified HHV-6 in 4/20 (20%) samples, using a combination of read numbers and genome coverage to infer positivity (33). Multiple considerations feed into the interpretation of NGS data (Fig 5). The four patients positive for HHV-6 by NGS in this cohort all had clinical syndromes that could be compatible with HHV-6 infection, but due to a limited dataset we cannot attribute causality. Interestingly, positive HHV-6 PCR from young children with severe respiratory tract infections suggest a potential pathological role of the virus that is not well described in this population to date. In future, in situations when HHV-6 detection is deemed significant, this could support the introduction of antiviral therapy, and/or reduce exposure to broad-spectrum antibiotics.

**Figure 5:**
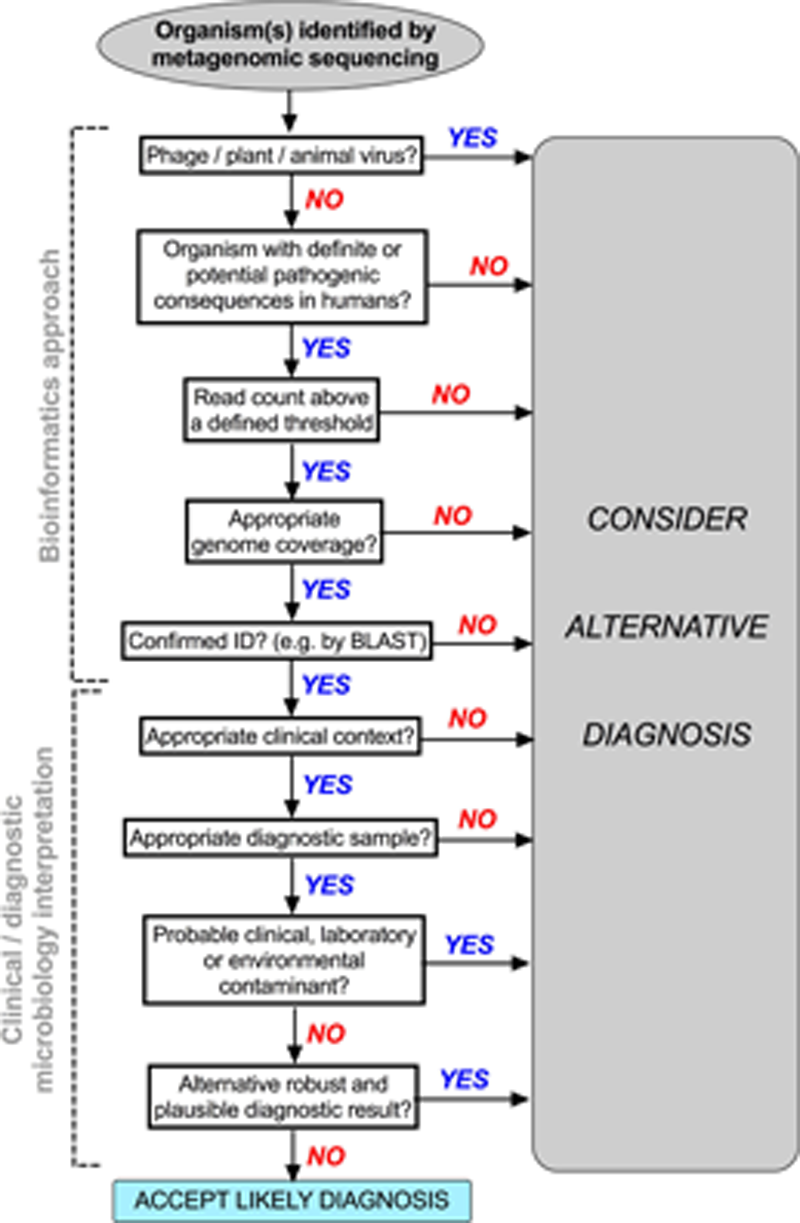
Suggested algorithm showing process of determining the significance of an organism identified from a clinical sample by next generation sequencing (NGS). This represents a structure that can be applied to bioinformatics and clinical interpretation of metagenomic data. We recognize that the approach and thresholds are different for different organisms, and that robust output also depends also on optimization of *in vitro* sample preparation.

The number of clinical requests for HHV-6 testing each year is small, even in a tertiary referral teaching hospital. By nature, our clinical sample set (HHV cohort) was strongly skewed towards sampling a high-risk population. Our data quality is dependent upon the completeness of electronic data, and it was not possible to determine temporality. For ongoing NGS work, priorities are screening blood samples, and development of positive internal controls to determining sensitivity. An international control standard for HHV-6 is currently being prepared by the National Institute for Biological Standards and Control (NIBSC).

Although it is doubtless at times a benign passenger, HHV-6 is indeed significantly associated with the clinical syndromes that arise in patients with profound immunocompromise in the haematology setting, and may have a role in other syndromes, including respiratory infections in children. Metagenomic approaches to clinical diagnostics are accelerating, and as additional data become available, increasing insights will be gained into the interpretation of these results.

## ACKNOWLEDGEMENTS

We are grateful to the Centre for Genomic Research (CGR), University of Liverpool, UK for running the HiSeq platform. Thank you to Dr Peter Muir (Public Health England South West Regional Laboratory, Southmead Hospital, Bristol), who provided valuable advice and discussion during data collection. A subset of these work were presented at the UK Federation of Infection Meeting, 2017 (34).

## AUTHORSHIP

Conceived and designed the experiments: CS, PK, PCM; Applied for ethics permission: PCM; Collected and curated clinical samples and data: MS, AM, NG, MA, KJ; Undertook laboratory work: CS, WFG; Analysed and presented the data: CS, TG, ALM, PCM; Wrote the article: CS, TG, ALM, DF, PCM, with feedback from all co-authors; Approved the final article: all authors.

## SUPPLEMENTARY RESOURCES

### Suppl data set 1. Human Herpes Virus 6 (HHV-6): a study of clinical laboratory data and next generation sequencing; https://doi.org/10.6084/m9.figshare.5671153.v1

This fileset includes the following:

- HHV6 cohort metadata (clinical cohort data to describe patients undergoing clinical testing for HHV-6 infection) as .xlsx and .csv files;
- VS cohort metadata (research cohort data to describe CSF and respiratory samples underdoing screening for HHV-6 infection using PCR and next generation sequencing) as .xlsx and .csv files.

